# High Prevalence of Strongyloidiasis in Spain: A Hospital-Based Study

**DOI:** 10.1101/852558

**Authors:** Ana Requena-Méndez, Joaquin Salas-Coronas, Fernando Salvador, Joan Gomez-Junyent, Judith Villar-Garcia, Miguel Santin, Carme Muñoz, Ana Gonzalez-Cordon, Maria Teresa Cabezas Fernández, Elena Sulleiro, Maria del Mar Arenas, Dolors Somoza, Jose Vazquez-Villegas, Begoña Treviño, Esperanza Rodríguez, Maria Eugenia Valls, Carme Subirá, Jose Muñoz, on behalf of the STRONG-SEMTSI working group

## Abstract

Strongyloidiasis is a prevailing helminth infection ubiquitous in tropical and subtropical areas. However, prevalence data are scarce in migrant populations.

This study aims at evaluating the prevalence of *S. stercoralis* at hospital level in migrant populations or long term travellers being attended in out-patient and in-patient units as part of a systematic screening implemented in 6 Spanish hospitals. A cross-sectional study was conducted and systematic screening for *S. stercoralis* infection using serological tests was offered to all eligible participants. The overall seroprevalence of *S. stercoralis* was 9.04% (95% confidence interval [95%CI] 7.76 −10.31). The seroprevalence of people with a risk of infection acquired in Africa and Latin America was 9.35% (95%CI 7.01-11.69), 9.22% (7.5-10.93), respectively. The number of individuals coming from Asian countries was significantly smaller and the overall prevalence in these countries was 2.9% (95%CI −0.3; −6.2). There was only one case (1/14 (7.14%) from an individual from East European countries. The seroprevalence in units attending potentially immunosuppressed patients was significantly lower (5.64%) compared with the seroprevalence in other units of the hospital (10.20%) or Tropical diseases units (13.33%) (p<0.001). Conclusions: We report a hospital-based systematic screening of strongyloidiasis with a seroprevalence of almost 10% in a mobile population coming from endemic areas suggesting the need of implementing strongyloidiasis screening in hospitalized patients coming from endemic areas, particularly if they are at risk of immunosuppression.

**Author summary:** Strongyloidiasis is an infection caused by the helminth *Strongyloides stercoralis* which is ubiquitous in tropical and subtropical areas. In the rest of the countries, it is also frequent in migrants coming from tropical and subtropical areas. The disease is more severe when an infected subject has an impaired immune system. Within this study we have evaluated the prevalence of this infection in people being attended in six Spanish hospitals. The prevalence was around 9%, being higher in Africa and Latin America compared with other regions. In addition, the prevalence in patients with an impaired immune system (immunosuppression) was lower compared with people non suffering immunosuppression. These results suggest that the prevalence of strongyloidiasis is quite high among migrants living in Spain and that a screening programme should be designed, particularly in immunosuppressed patients that are at more risk of suffering severe complications of the infection.

## INTRODUCTION

Migration flows from American, African and Asian countries to Europe have shown that a high percentage of arriving individuals may be chronically infected with *S. stercoralis*, what may have a public health impact in non-endemic countries that are hosting these populations[1, 2]. The infection is ubiquitous in tropical and subtropical areas although it may also occur in temperate countries with the appropriate conditions such as certain areas of Spain or Italy [3–5]. Worldwide estimates based on standard faecal techniques have suggested that between 30 and 100 million people worldwide are infected. These figures may be underestimated due to the low sensitivity of the traditional diagnostic methods [1, 6].

Unlike other parasitic infections, this helminth has some characteristics that are of particular importance for migrant populations [7]. In the first place, the infection can persist the whole lifetime due to its ability to replicate in the human host [8]. Therefore, people coming from endemic areas may be at risk their whole life irrespective of the moment they arrive to a non-endemic area as long as they are not treated. Second, *Strongyloides stercoralis* infection is generally asymptomatic or causes unspecific symptoms, being unnoticed for health professionals not looking for it [9]. Third, although the infection is rarely transmitted through person to person, [10] it can be transmitted through organ transplantation, and autochthonous cases have been reported in non-endemic areas [11]. Thus, screening should be considered in potential donors at risk of the infection [12–14]. Finally, in case of immunosuppression, particularly described with the concomitant use of steroids, transplant recipients or patients with malignancies and Human T-Cell Lymphotropic virus-1 co-infections, the parasite may enter into a high replicating cycle (called hyperinfection) or disseminate to vital organs (disseminated strongyloidiasis), causing a severe disease with high mortality [15]. In this regard, immunosuppression permit larval proliferation (hyperinfection disease) and also dissemination to other organs (disseminated *Strongyloides*) [15]. The diagnosis of strongyloidiasis in non-endemic areas where re-infection is unlikely is currently based on a serological test which has considerably higher sensitivity compared with standard faecal techniques [8]. Despite having cross-reactions with other helminthic infections, this fact is less likely to occur in migrant populations since the possibility of co-infections is lower [16] and therefore it is nowadays the current recommended screening technique in these populations[17]. The sensitivity of the serological tests in immunosuppressed individuals seems to be lower [18]; but these are limited data and further prospective studies should better evaluate the accuracy of the serological tests in immunosuppressed patients.

Even with the limitations of current diagnostic methods, screening of high-risk groups and treatment of infected individuals is of key importance. In this regard, screening of strongyloidiasis in newly arrived migrants has been recommended by the European Centre for Disease Prevention and Control [19, 20], particularly in immunosuppressed individuals, given the potential individual morbidity and mortality [16].

Evidence of prevalence of strongyloidiasis in migrant populations is scarce although it is known to vary substantially depending on the country of origin, being particularly high in countries such as Cambodia (36%) or Latin American countries (26%) [21]. Hospital-based prevalence studies conducted all of them in specialized units suggest prevalence of *S. stercoralis* between 4.5 and 11% in migrant populations [22–24].

Available data suggests a higher prevalence of the infection in immunosuppressed individuals at risk [25–27].

Our study aims at evaluating the prevalence of *S. stercoralis* at hospital level in migrant populations or long term travellers being attended in out-patient and in-patient units as part of the systematic screening implemented in 6 Spanish hospitals.

## METHODS

### Study population, data collection and patient management

This is a cross-sectional study, carried out as part of a hospital-based prospective cohort study conducted in 6 referent hospitals in Spain (Hospital Clinic, Hospital del Mar, Hospital Universitari Bellvitge, Hospital Sant Pau and Hospital Vall d’Hebron located in Barcelona and Hospital de Poniente in Almeria province) aimed to evaluate the prevalence and risk factors associated with *Strongyloides stercoralis* infection, particularly related to immunosuppressed patients.

Although a systematic screening has not been widely implemented at national level, it was established that individuals either hospitalized or being attended in any specialized out-patient or in-patient unit of the hospital should be screened for strongyloidiasis. Not all hospital units or department were participating in the study. In particular, infectious diseases or Tropical unit and units attending severe immunosuppressed patients were willing to participate and to implement the systematic screening.

Migrant adult individuals or adult long-term travellers (defined as those staying one year or more in any endemic country) were systematically offered being screened for *Strongyloides stercoralis* based on a serological test. Endemic countries were considered any country in Asia, Oceania, Africa, East European countries or Latin American countries. Therefore, inclusion criteria were to have reside in an endemic country for one more than year and to be attended in the hospital for any reason and irrespective of the level of immunosuppression.

As part of this programme, all individuals who met inclusion criteria for screening were offered to be tested for *S. stercoralis* infection through a serological blood test.

All positive patients were invited to participate and signed the consent form and were provided treatment with ivermectin 200 mcg/kg during 2 consecutive days. Participants were clinically and microbiologically followed – up after six months of therapy as part of the routine clinical practice.

### Microbiological procedures serology of *S. stercoralis*

Serum samples were tested for specific antibodies using a commercial Enzyme Linked Immunosorbent assay (ELISA) assays according to the manufacturer’s instructions. A single ELISA test was used (Strongyloidiasis ELISA Kit based on IVD *Strongyloides stercoralis* crude antigen). To reduce the inter-laboratory variability, an internal workshop was carried out before starting the study to standardized the serological techniques and also the interpretation of the results. Positive samples are defined by absorbance greater than 0.2 OD units. The absorbance of study sample/0.2>=1 (calculated value) was used as the cut-off.

### Sample size estimation

Based on other seroprevalence studies conducted in non-endemic areas, we estimated a prevalence of *S. stercoralis* of 11% for our cohort [17]. Based on these data, it was estimated to screen 1700 individuals during 24 months to recruit 150 cases of strongyloidiasis, considering an Alfa error of 5% and a lost to follow-up of 10%.

### Statistical analysis

Categorical variables were described using a frequency distribution and the median and the interquartile range were used to describe age. Countries of origin were subsequently grouped into different geographic areas according to the criteria that could more precisely identify the migration patterns. The GeoSentinel geographic area distribution was selected [28]. Prevalence point estimates and their 95% confidence intervals (95%CI) were obtained. We used Fisher’s exact test to compare percentages. Data were managed and analysed using STATA 13. We built a world map with country of acquisition’s prevalence using the free software QGIS, version 2.18.22.

Hospital Units identified as attending potentially immunosuppressed patients were: Haematology, Oncology, HIV, Autoimmune Diseases, Rheumatology, Transplant Units and Oncology.

#### Ethics

This project was part of a study where, individuals with a positive for *S. stercoralis* test were further invited to participate to evaluate risk factors associated with the infections particularly related to immunosuppression and signed a written consent form. The study was approved by the ethic committee of all hospitals participating in the study (Hospital Clinic HCB/2014/0321). Only adult individuals participated in this study.

The study has been reported following the STROBE statement (Strengthening the reporting of observational studies in epidemiology) for reporting observational studies (Annex 1).

## RESULTS

From October 2014 to September 2016, 2,024 individuals attended in the 6 hospitals were screened for strongyloidiasis and 77 of them were excluded from the study because either the screening had not been done with a serological test or it could not be assured a systematic approach in several units of one hospital. Thus, 1,948 individuals were finally included. Basic demographic data are summarized in table 1. The median age was 38 years (Interquartile range 31-47) and 57.96% were male; 804 (41.27%) were screened at Infectious diseases or Tropical diseases units, 709 (36.40%) in HIV-units, 150 (7.70%) in transplant units, 101 (5.18%) in Oncology or Haematology units, and 86 (4.41%) in Rheumatology or Autoimmune diseases departments or other units addressing specifically immunosuppressed patients. Finally, 77 (3.95%) and 21 (1.08%) were recruited in general services of the hospital (e.g. Emergency, Internal Medicine) and in 21 (1.08%), the origin-department was unknown.

**Table 1.** General characteristics of individuals screened for strongyloidiasis

|  |  |
| --- | --- |
| **Age (n=1943)** |  |
| Median (IQR) | 38 (31-48) |
| Group |  |
| <25 | 201 (10.34%) |
| 26-39 | 846 (43.54%) |
| 40-54 | 652 (33.56%) |
| >54 | 244 (12.56%) |
| **Male sex (n=1634)** | 947 (57.96%) |
| **Hospitals** |  |
| Clinic | 902 |
| Vall d'Hebron | 411 |
| Hospital Poniente | 303 |
| Hospital Mar | 198 |
| Hospital Sant Pau | 70 |
| Hospital Bellvitge | 64 |
| **Departments or units** |  |
| General services | 77 |
| Autoimmune/Rheumatology | 86 |
| Transplant units | 150 |
| Haemato-Oncology | 101 |
| International Health | 804 |
| HIV units | 709 |
| Other | 21 |
| **Continents (n=1876)** |  |
| **Africa** | 600 |
| North Africa | 141 |
| SSA | 459 |
| **America** | 1106 |
| South America | 934 |
| Central America & Caribe | 172 |
| **Asia** | 103 |
| South - Central Asia | 61 |
| South-East Asia | 27 |
| Middle East | 3 |
| East Asia | 12 |
| **Europe** | 67 |
| West Europe | 53 |
| East Europe | 14 |

Of the total screened population, 58.96% (1106/1,876) had a Latin-American origin, 31.98% (600/1,876) were from Africa, 5.49 (103) from Asia and 3.57% (67) from Europe.

There were differences across the centres concerning the country of origin of individuals screened. This could be explained by the demographics of the migration flows in Spain: In the hospital located in Almeria, which is very close to the African border, 94.39% had an African origin whereas for the rest of hospitals, more than half of individuals have a Latin-American origin (table 1). Services/units of hospitals were also grouped into those units attending essentially immunosuppressed or potentially immunosuppressed patients (HIV, transplant, haematology, oncology, autoimmune diseases units) compared with units that do not attend immunosuppressed patients (Infectious diseases, Tropical diseases units or general services of the hospital). Latin American migrants were slightly more frequent in the potentially immunosuppressed group (67.19%) compared with the other (49.12%) (P-value <0.001).

### Seroprevalence

From 1947 patients tested, 176 had a positive serological test for *S. stercoralis*. The overall seroprevalence of *S. stercoralis* was 9.04% (95%CI 7.76 −10.31). The prevalence varied significantly among the centres from 6.57% (13/198) in Hospital del Mar to 29.69% (19/64) in Hospital de Bellvitge (p<0.001).

Women had a higher seroprevalence (10.77%) compared with men (6.96%) (p-value = 0.005). We found no association between the age group and the prevalence (p= 0.962).

### Seroprevalence by geographic distribution

The seroprevalence of people with a risk of infection acquired in Africa was 9.35 (95%CI 7.01-11.69), being in Sub-Saharan Africa (SSA) countries higher (10.89%; 95%CI 8.03-12.75) compared with North African countries (4.29%, 95%CI 0.8-7.69), (p= 0.019).

Morocco was the country of acquisition from Africa with a larger population screened, showing a seroprevalence of 4.69% (95%CI 0.9-8.34). There were only 8 countries with >= 20 observations (See figure 1) and the results of prevalence by country of origin are summarized in Annex 2.

**Figure 1.**
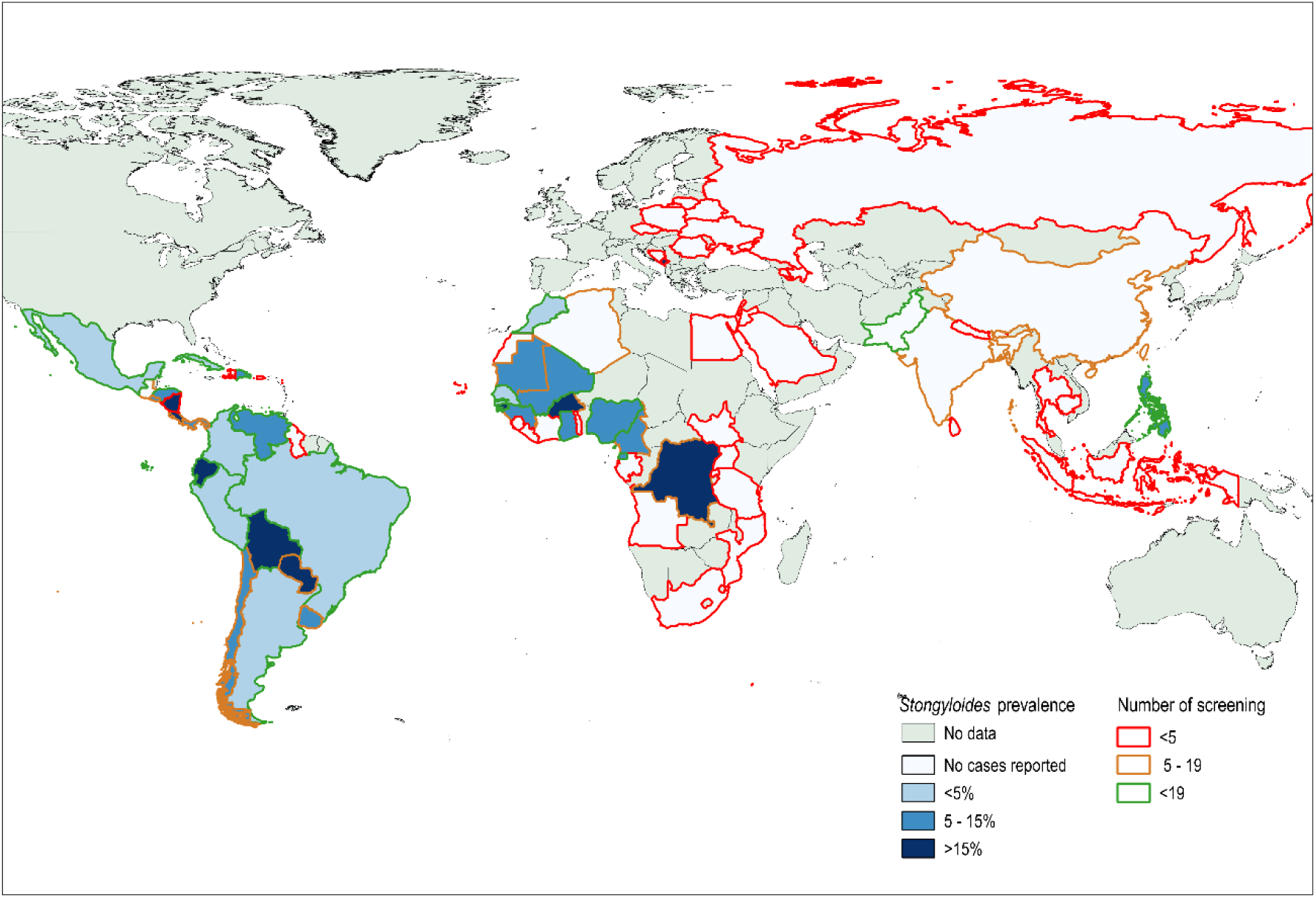
Seroprevalence of strongyloidiasis in migrants and travellers in Spain by country of acquisition Legend: The figure was built with the QGIS software, version 2.18.22

The Latin America (LA) continent showed a seroprevalence of 9.22% (95%CI 7.5-10.93), being the seroprevalence in people coming from South American countries 9.64% (95%CI 7.74-11.5) and 6.98% (95%CI 3.13-10.82) the prevalence in people coming from Central America and the Caribbean (p=0.268). Ecuador and Bolivia were the countries of acquisition with the highest seroprevalence, 17.48% (95%CI 10.01-24.9) and 15.81% (95%CI 11.84-19.77), respectively. The prevalence was below 8% in the rest of the countries with more than 50 people screened. (see figure 1)

The number of individuals screened coming from Asian countries was significantly smaller and the overall prevalence in these countries was 2.9% (95%CI −0.3, 6.2). Only 3 cases were observed, in people coming from South-East Asia (11.11%; 95%CI 1.5-23.78) and all 3 were originally from Philippines (seroprevalence 13.63%).

There was only one case (1/14 (7.14%) from an individual from East European countries. The patient was from Montenegro and he had not travelled outside Europe.

There were 50 individuals originally from West Europe that were also screened for *S.stercoralis* after long term travels to endemic countries. In the 3 out of 50 individuals (5.6%) with a positive result, the country of acquisition could not be identified since all of them had undertaken trips to several endemic areas in the past (Latin America and South East Asia countries).

### Seroprevalence by screening department

The seroprevalence in units attending immunosuppressed and potentially immunosuppressed patients was significantly lower (5.64%) compared with the seroprevalence in other units of the hospital (10.20%) or Tropical diseases units (13.33%) (p<0.001) (See table 2). HIV and transplants units showed the lowest prevalence rates (4.51% and 2% respectively).

**Table 2.** Seroprevalence by geographic area (GeoSentinel areas)

|  | Percentage (CI) | A: Immunosuppressed units* (%) | B: Other Non immunosuppressed units from the hospital ** (%) | C: Tropical diseases units | p-value*** |
| --- | --- | --- | --- | --- | --- |
| South-America | 9.6 (7.8-11.7) | 30/558 (5.38%) | 1/11 (9.1) | 59/365 (16.2%) | <0.001 |
| Central America and Caribbean | 7 (3.7-11.9) | 5/128 (3.91) | 0/8 (0) | 7/36 (19.44) | 0.04 |
| North African | 4.3 (1.6-9.1) | 3/90 (3.3) | 0/3 | 3/47 (6.4) | 0.658 |
| Sub-Saharan Africa | 10.9 (8.2-14.1) | 12/155 (7.7) | 1/6 (16.7) | 37/298 (12.4) | 0.286 |
| Middle East | 0 (0-70.6) | 0/2 | 0 | 0 | --- |
| South-Central Asia | 0 (0-58.7) | 0/29 | 0/5 | 0/27 |  |
| South-East Asia | 11.1 (2.4-29.2) | 2/19 (10.5) | 0/2 | 1/6 (16.7) | 0.801 |
| East Asia | 0 (0-26.4) | 0/5 | 0 | 0/7 | --- |
| East Europe | 7.1 (0.2-33.9) | 1/13 (7.7) | 0/1 | 0 | 0.773 |
| Western Europe (travellers with multiple travels) | 5.7 (1.2-15.7) | 3/21 (14.3) | 0/15 | 0/17 | 0.089 |
| Total | 9 (7.7-10.3) | 59/1046 (5.64) | 10/98 (10.2) | 107/803 (13.39) | <0.001 |
| *Immunosuppressed units: Haematology, Oncology, Autoimmune diseases, Rheumatology, HIV and Transplant Units; **Other non-immunosuppressed units: Internal Medicine, Emergency, Cardiology, Neurology, Dermatology and Pneumology; Chi square test used to calculate difference in prevalence in units A, B and C |  |  |  |  |  |

In these units, the rate in individual with a LA origin was 3.74% and 3.39% respectively, significantly lower compared with other units.

## Discussion

Our study shows an overall prevalence of almost 10% which is consistent with other studies conducted in migrant population living in non-endemic areas [17].

This is one of the first studies evaluating the results of a systematic screening in potentially immunosuppressed patients in which the disease may be more severe.

The high prevalence found in our study supports the need of screening strategies in patients that are potentially immunosuppressed particularly for Sub-Saharan African and Latin American migrants.

Potentially immunosuppressed people are at higher risk of developing a severe complication of the disease as it has been extensively reported elsewhere[15]. However, the probability of developing hyperinfection or dissemination of the infection in the high-risk immunosuppressed population is unclear. Systematic screening has been recently recommended in high-risk population coming from endemic areas [19]. However, few programmes have reported their results so far. In this regard, a reference centre in Austria has initiated screening of *S.stercoralis* to all transplant recipients showing 3% prevalence even though only a small percentage were migrants and most of them from East European countries [29].

Interestingly, the seroprevalence in Ecuador and also in Bolivia was much higher compared with other LA countries but similar compared with other studies conducted in migrants from these countries [30], which may be partially explained due the profile of migrants coming from rural areas in these countries. In addition, the sampling in these countries may have been also overrepresented compared to other LA countries and therefore the prevalence in LA may be overestimated.

In other geographic areas, such South-East Asia or North Africa, our data show a not negligible rate with seroprevalence higher than 5% in countries such as Morocco or Philippines. We have not found any other published seroprevalence data in such countries. Although the limited sampling may prevent to extract conclusion about the real prevalence in these countries, these data refute the need to consider them in the pool of countries where screening should be recommended.

Data from Asian countries are still very scarce and the limited sampling may prevent to extract proper conclusions regarding the prevalence in this geographic area. Although we have not found studies providing specifically prevalence estimates in Asian migrants, some surveillance data about health conditions in migrants have reported a 5-7% of strongyloidiasis among all diagnosis in people coming from South-East Asia and South Asian countries[31]. Further studies in non-endemic areas should better evaluate the *S. stercoralis* prevalence in people coming from Asian countries.

Seroprevalence in women was found to be significantly higher compared to men which differ from other studies where the prevalence was more frequent in males [32]. In our population data-base some countries like Bolivia with a high seroprevalence have a larger proportion of women. This fact may have partially explained this difference.

In our study, prevalence in HIV-infected individuals and transplant units was much lower compared with other general out/in-patients units services of the hospitals. One possible explanation that could have contributed to the lower prevalence found in potentially immunosuppressed patients, is that the serology has a lower sensitivity in immunosuppressed patients as it has been reported in other studies [18].The sensitivity of the serology in immunosuppressed patients deserves further evaluation of its accuracy, since only few studies have evaluated it, showing a lower sensitivity compared to parasitological techniques [18] and no studies have evaluated its accuracy at different levels of immunosuppression. In addition, it is still not clear if the serology should not be recommended in such cases or if the screening should be performed adding other direct parasitological techniques [16].

Unfortunately, and due to the study design, we could not have the information about the level of immunosuppression that patients had at the time the screening test was performed. Therefore, we cannot estimate to what extent the lower-sensitivity of the screening test is causing this difference. Further prospective studies should better evaluate the accuracy of the serological test in immunosuppressed patients including the assessment of the accuracy with different levels or categories of immunosuppression.

It must be added that the prevalence of Latin-American migrants screened in these units (HIV and transplant) was particularly lower compared with the rest of units of screening. The different profile of Latin American migrants attended in the HIV and transplant units compared with Tropical units (e.g. migrant origin from urban vs. rural areas) could partially explain it.

This is a hospital-based study and results should be interpreted in the context of a hospital-based population. Generalizing our prevalence results to the general (or wider) population should be done with caution since the positive selection bias should be taken into account. In addition, another limitation is the difference of the prevalence between the hospitals participating in this study that could be partially explained by differences in the migrant profile. However, a lack of systematic screening could have overestimated the prevalence in some cases.

### Conclusions

We report a hospital-based systematic screening of *Strongyloides* with a seroprevalence of almost 10% in migrant population from endemic areas what put in evidence the need of implementing strongyloidiasis screening strategies in hospitalized patients, particularly if they are at more risk of immunosuppression.

## Acknowledgements

This work was funded by the Spanish Society of Tropical Medicine and International Health. A. ISGlobal is partially supported by the RICET (RICET (RD12/0018/0010) within the Spanish National plan of R þ D þ I, which is co-funded by ISCIII-(FEDER)).

## Financial disclosure form

The funders had no role in study design, data collection and analysis, decision to publish, or preparation of the manuscript.

## Supporting information legends

Supplementary file: STROBE statement - Checklist of items that should be included in reports of cross-sectional studies

